# *eyeris:* A flexible, extensible, and reproducible pupillometry preprocessing framework in R

**DOI:** 10.1101/2025.06.01.657312

**Authors:** Shawn T. Schwartz, Haopei Yang, Alice M. Xue, Mingjian He

**Author notes:** For correspondence (S.T.S.).

## Abstract

Pupillometry provides a non-invasive window into the mind and brain, particularly as a psychophysiological readout of autonomic and cognitive processes like arousal, attention, stress, and emotional states. Pupillometry research lacks a robust, standardized framework for data preprocessing, whereas in functional magnetic resonance imaging and electroencephalography, researchers have converged on tools such as *fMRIPrep*, *EEGLAB* and *MNE-Python*. Many established pupillometry preprocessing packages and workflows fall short of serving the goal of enhancing reproducibility, especially since most existing solutions lack designs based on Findability, Accessibility, Interoperability, and Reusability (FAIR) principles. To promote FAIR and open science practices for pupillometry research, we developed *eyeris*, a complete pupillometry preprocessing suite designed to be intuitive, modular, performant, and extensible (https://github.com/shawntz/eyeris). Out-of-the-box, *eyeris* provides a recommended preprocessing workflow and considers signal processing best practices for tonic and phasic pupillometry. Moreover, *eyeris* further enables open and reproducible science workflows, as well as quality control workflows by following a well-established file management schema and generating interactive output reports for both record keeping/sharing and quality assurance of preprocessed pupil data prior to formal analysis. Taken together, *eyeris* provides a robust all-in-one transparent and adaptive solution for high-fidelity pupillometry preprocessing with the aim of further improving reproducibility in pupillometry research.

## 1 Introduction

The measurement of pupil size and its relation to cognition began more than a century ago ***(Haab, 1886)***. Our current understanding of pupil size is that it is an integrated psychophysiological readout of cognitive processes. As such, pupillometry offers a non-invasive window into the human mind and brain.

The breadth of cognitive processes that have been associated with pupil size is expansive, including, but not limited to, emotional arousal ***(Bradley et al., 2008; Oliva and Anikin, 2018)***, mental effort ***(van der Wel & van Steenbergen, 2018; Kahneman and Beatty, 1966; Kramer et al., 2013)***, fatigue/task performance ***(Aston-Jones and Cohen, 2005; Eldar et al., 2013; Knapen et al., 2016; McGinley et al., 2015; Murphy et al., 2011)***, response preparation ***(Einhäuser et al., 2008; Reimer et al., 2014)***, uncertainty/decision-making ***(Urai et al., 2017; Gee et al., 2014; Jepma and Nieuwenhuis, 2011; Kawaguchi et al., 2018)***, prediction error ***(Alamia et al., 2019)***, working memory ***(Keene et al., 2022; Robison and Unsworth, 2019; Unsworth and Robison, 2015, 2017a,b)***, and episodic memory (*C****lewett et al., 2020; Madore et al., 2020; Madore and Wagner, 2022; Miller et al., 2019; Miller and Unsworth, 2020; Robison et al., 2022; Schwartz et al., 2025)***.^1^

Pupil size has also been linked to activity of specific neural substrates, for example, pupil diameter dynamics are oftentimes interpreted as a non-invasive (indirect) surrogate of locus coeruleus-noradrenergic (LC–NA) neuromodulatory activity ***(He et al., 2020; Joshi et al., 2016; Joshi & Gold, 2020; Murphy et al., 2014; cf. Megemont et al., 2022)***. Additionally, it is worth noting that given LC’s central role in regulating autonomic function and arousal ***(Mather & Harley, 2016; Poe et al., 2020; Samuels and Szabadi, 2008)*** and that LC is the first brain region to develop tau pathology in Alzheimer’s disease ***(Betts et al., 2019; Braak and Del Tredici, 2011; Holper et al., 2022; Spires-Jones and Hyman, 2014)***, pupillometry is also being used as a noninvasive window into cognitive aging ***(Hämmerer et al., 2018; El Haj, 2024; Kim et al., 2024; Piquado et al., 2010)***.

Consequently, collecting high-quality pupil data and appropriately analyzing these data are critical for advancing both theories of cognition as well as translational applications of pupillometry. Moreover, data preprocessing is a fundamental prerequisite to high quality pupillometry data analysis ***(Botvinik-Nezer et al., 2020; Fink et al., 2024; Hershman et al., 2024; Kret and Sjak-Shie, 2019; Laeng and Mathôt, 2024; Mathôt and Vilotijević, 2023; Steinhauer et al., 2022; van Rij et al., 2019)***.^2^ In pupillometry research, experimenters deal with fewer time series, at most two for binocular recordings, compared to electroencephalography (EEG) or fMRI, but many of the same principles apply: handling missing data, excluding artifacts, and smoothing out physiologically irrelevant noise, to name a few.

Numerous researchers have contributed excellent software packages to the community over the years to aid others in preparing and analyzing pupillometry data without the need to implement core preprocessing functions themselves (e.g., ***Forbes, 2020; Geller et al., 2020; Hershman et al., 2019; Kinley and Levy, 2021; Tsukahara, 2020)***. These packages vary in how much preprocessing flexibility they afford: several expose adjustable parameters and modular, user-configurable workflows (e.g., ***Geller et al., 2020; Tsukahara, 2020)***. However, a standardized preprocessing pipeline with reproducibility baked into its design remains a critical gap in current pupillometry research, as noted by ***Mathôt and Vilotijević (2023)***:

There is no commonly agreed-upon workflow for conducting cognitive-pupillometry experiments: design criteria, preprocessing steps, and statistical analyses all differ vastly between studies. Some attempts at standardization have been made, but these are not easily applicable to cognitive pupillometry […] Furthermore, guidelines are often conceptual rather than specific implementations, and the lack of concrete examples and code makes it difficult for researchers to incorporate these guidelines into their own workflow. (p. 3055)

With reproducibility as a core design philosophy, we built eyeris, an R package dedicated to providing unified standards for pupillometry preprocessing ***(Schwartz, 2025a)*** that improves on where current tools often fall short: the transparency and shareability of the preprocessing pipeline as a whole. While *eyeris* is not qualitatively more prescriptive than existing tools, it offers a standardized, out-of-the-box means of documenting and exporting the complete preprocessing record in a form ready to be published alongside a research study.

With *eyeris*, we not only intend to provide users with an extensible toolkit for conducting robust pupillometry preprocessing, but critically aim to contribute to standardizing best practices in pupillometry research (especially for cognitive pupillometry; see ***Mathôt (2018a)*** for a review discussing pupil responses, the neural pathways subserving these pupillary responses, and their relationship to high-level cognition). We should note that we are not the first group to put forth a recommended set of preprocessing guidelines for cognitive pupillometry. Here, we have built upon the comprehensive guide to cognitive pupillometry preprocessing put forth by ***Mathôt and Vilotijević (2023)***; their stated aim is to provide “guidelines [that] are meant to be illustrative, rather than prescriptive; that is, we show how things could be done (rather than how they should be done) in order to implement a workflow that is both easy to implement and that meets contemporary standards for good scientific practice” (p. 3056). In contrast, we adopt a *guided and reproducible* rather than merely *illustrative* approach to pupillometry preprocessing workflows. We did not take a strictly prescriptive approach as we recognize that our knowledge on all possible experimental designs involving pupillometry is too limited to prescribe a set of universally optimal parameters. At the same time, from good practices in general signal processing and existing domain expertise in pupillometry research, we are confident that *eyeris* can provide more guidance than existing toolkits to users so that they can avoid preprocessing choices that are likely suboptimal for most investigations of pupil size. To illustrate, some tools suboptimally apply core operations such as interpolation and smoothing to already-epoched, trial-level segments rather than to the continuous recording, which can introduce discontinuities at trial boundaries; *eyeris* performs these operations on the whole timeseries prior to epoching (see ***Section 2.4.1)***.

Without consistent preprocessing, studies leveraging pupillometry to advance key theories in psychological science could be hindered by high degrees of freedom and vulnerable to unintended mistakes during preprocessing ***(Aczel et al., 2026; Steegen et al., 2016)***. As such, our accompanying R package is designed with “glass box” principles (see ***Esteban et al., 2019; Poldrack et al., 2019)***, thereby equipping both novice and experienced pupillometry researchers with key tools needed to appropriately transform raw pupillometry data into a format well-suited for downstream analysis, visualization, and statistical inference. Embracing a carefully designed pipeline is highly beneficial, as it enables researchers to focus on hypothesis testing with minimal worry about whether the data were preprocessed correctly. In this era of open-source software, it is also important to allow experienced users to inspect and modify steps in a preprocessing pipeline with minimal hurdles.

*eyeris* strongly encourages users to follow a standard set of steps in a particular order and with carefully selected default parameters. The overarching aim is to avoid unnecessary preprocessing decisions that may lead to issues when made with incomplete knowledge of some of the nuances of signal processing. At the same time, *eyeris* provides transparent access and configurability to all preprocessing steps in a user-friendly fashion, which is supported by its modular, glass-box design (***Figure 1***, ***Box 1)***. This design approach greatly reduces the burden of usage by removing the need to develop lengthy scripts for individual use cases ***(Varoquaux, 2016)***.

**Figure 1.**
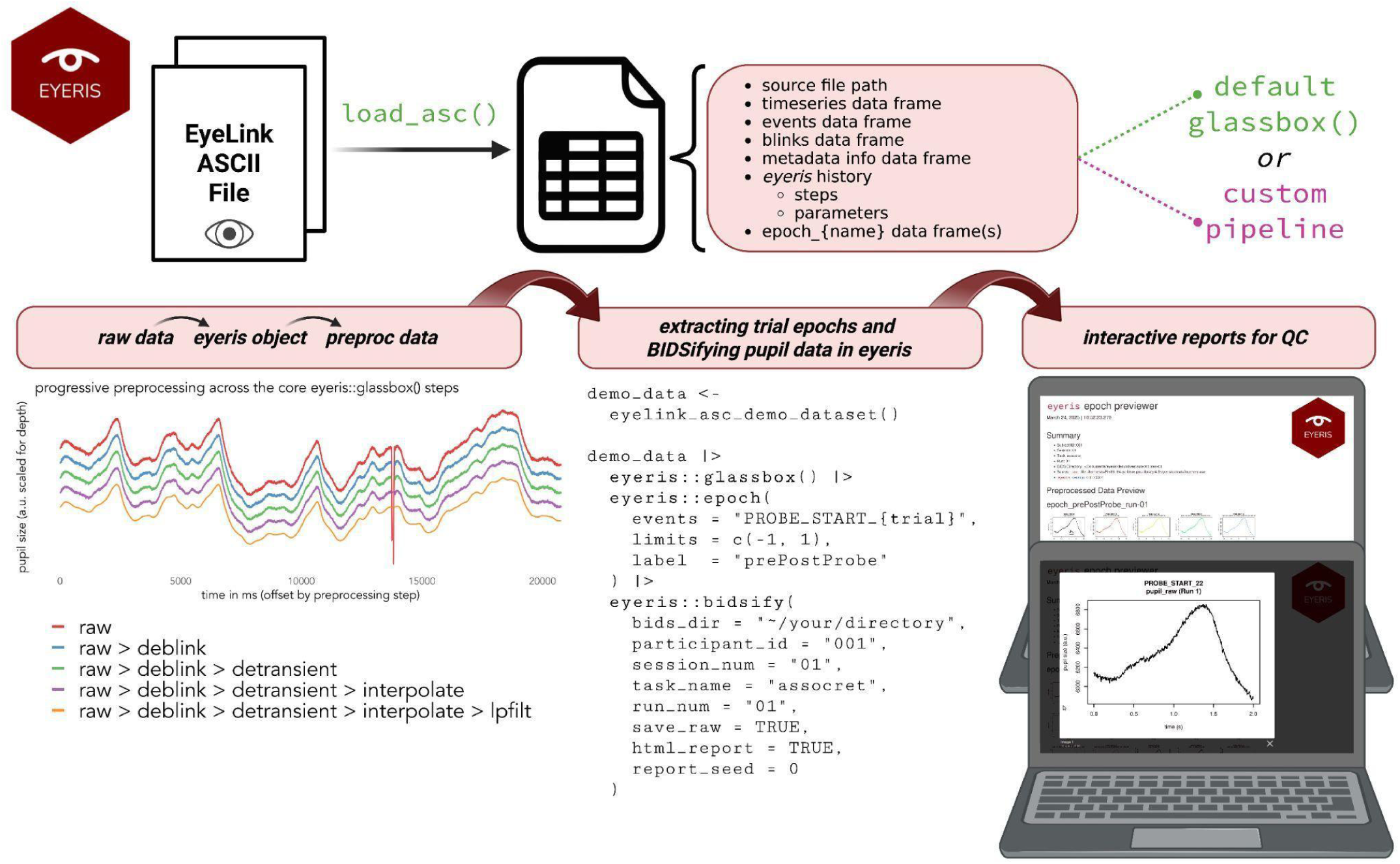
A sample workflow for the *eyeris* R package. The example code shows an end-to-end preprocessing pipeline, starting at raw data from the tracker to epoched and derived files ready for visualization and analysis. Here, we draw particular attention to the flexibility *eyeris* offers for segmenting (i.e., epoching) trials from the continuous pupil time series. In this demonstration, the *epoch() function* will find all event messages from the recording session that include the “PROBE_START_*” wildcard expression and extract 1-second prior to as well as 1-second following the onset timestamp from the matched message string (and will name the resulting tidy data frame of extracted pupil epochs: “prePostProbe”, which will be saved to the resulting *eyeris* S3 class “list” object — *not shown here*). Note, the “{trial}” syntax is a feature that enables a user to tell *eyeris* to not only treat that portion of the string as a wildcard expression, *but also* to take whatever substring matched to that position of the string and place its content in a new column with the header title: *trial*. This feature was designed with the intention to streamline the extraction of metadata embedded within tracker event messages into the same tidy data frame as any corresponding pupil data extracted from these events. The *epoch()* function is quite flexible and can handle a wide range of event message-string configurations; an in depth walkthrough can be found in the package documentation. Created in BioRender. *Schwartz (2025b)* https://BioRender.com/vkahl3u.

## 2 Method

Pupillometry time series require a relatively small number of preprocessing steps compared to multivariate data (such as EEG and fMRI); yet, there remains a defined set of issues to be addressed when preprocessing pupil size data. Critically, these preprocessing steps must be completed in a certain order to avoid unexpected modifications of the data in a way that could compromise downstream analyses (see ***Šoškić et al., 2024)***. In this section, we describe some of the most common preprocessing steps for pupillometry data, which are implemented in the current version of *eyeris*. While additional steps might be added in the future, some of the basic principles covered here will be applicable in general.

### 2.1 eyeris R package

The *eyeris* R package is actively maintained and available on both CRAN and GitHub (see *Code Availability*). ***Box 2*** describes the three recommended methods for installing *eyeris*.

##### Box 1. Example usage of the *glassbox* pipeline wrapper & deconstructing *glassbox()*: using the flexible *eyeris* primitives

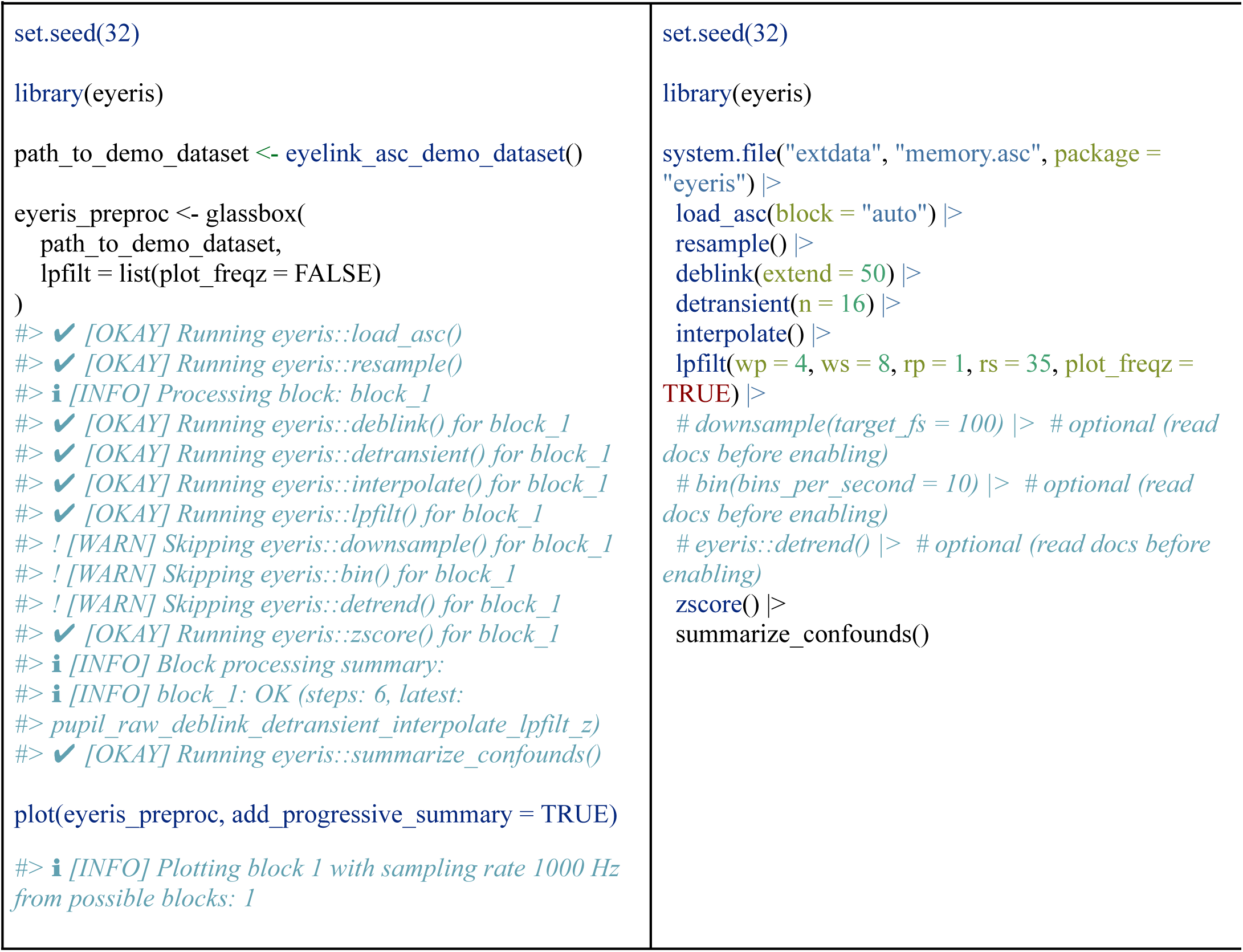

##### Box 2. *eyeris* R package installation options

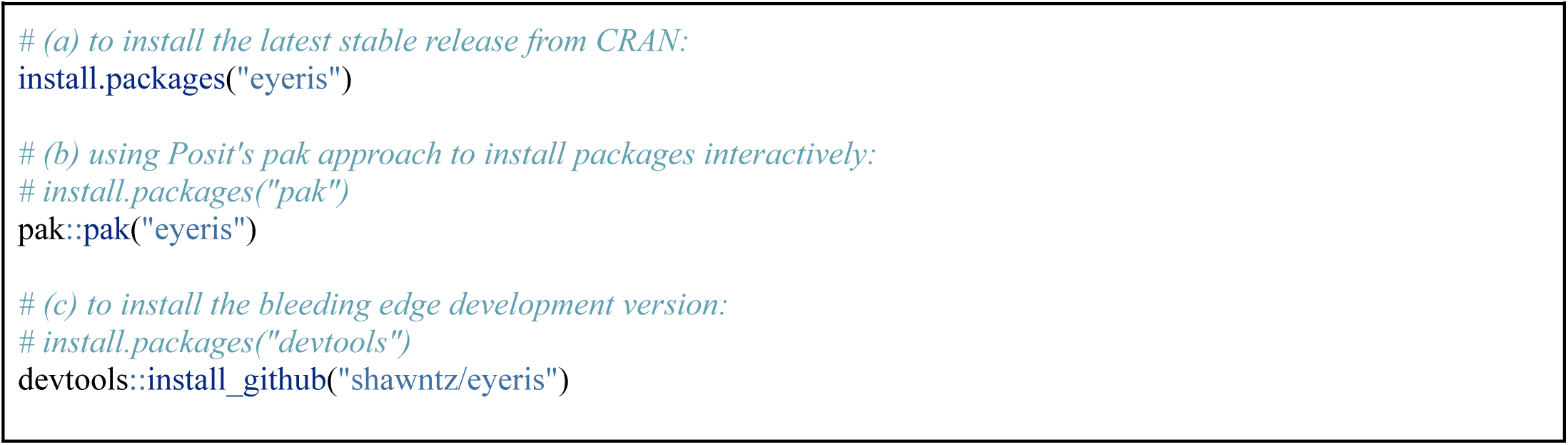

### 2.2 Entry-point function: glassbox()

The primary entry point for preprocessing pupillometry data in *eyeris* is the glassbox() function, which offers an opinionated, fully transparent preprocessing pipeline. This “glass-box” design contrasts with “black-box” workflows by allowing (and encouraging) users to understand and evaluate the consequences of each processing step on their data. Within *eyeris*, the glassbox() function completely streamlines the preprocessing workflow, enabling researchers to seamlessly transform raw pupil data into a high-quality, standardized format ready for analysis and visualization.

We designed glassbox() to operate in both *interactive* and *headless* modes. In the interactive mode, users are well-positioned to evaluate pre- and post-processing plots of their pupil time series data following the execution of each pipeline step, with prompts to continue or modify parameters as needed. In the headless mode, users are permitted to automate preprocessing of pupil datasets of any size, including large-scale pupillometry datasets on cloud computing clusters^3^, using recommended default parameters without manual intervention.

Users are permitted to override the default parameters of any preprocessing step within glassbox() by passing a named list of arguments to the specific function (e.g., deblink = list(extend = 100)) as arguments to the glassbox() function. Taken together, *eyeris*’ flexibility out-of-the-box facilitates real-time parameter optimization and empowers researchers to tailor the pipeline to their specific data characteristics and research questions in a transparent fashion.

### 2.3 Sequential preprocessing steps of the glass-box pipeline implementation and usage

The glassbox() pipeline executes a fully transparent *guided and reproducible* sequence of pupillometry preprocessing steps, which are also available as individual modular functions (see ***Box 1)***. Although users are able to build their own pipelines using the individual modules, we strongly recommend that users invoke the glassbox() pipeline function, given each step is critical and must be performed in a specific order to avoid unexpected data modifications that could compromise downstream analyses. Below, we demonstrate the sequence of core methods that comprise the glassbox() function to showcase both its flexibility and its syntactic simplicity.

#### 2.3.1 Loading and parsing data: eyeris::load_asc(), eyeris::resample()

This initial step builds upon the *eyelinker::read.asc()* function and parses SR Research EyeLink *.asc* files^4^, extracting raw time series data, event messages, blink events, and various metadata from the file header. The load_asc() function within *eyeris* also supports automatic handling of multiple recording segments (i.e., blocks) within a single .asc file and offers options for gracefully handling binocular data out-of-the-box (e.g., averaging left and right eye pupil sizes, or processing them independently). Note, it is critical that users load their data using eyeris::load_asc() given there are a number of configuration steps *eyeris* performs to ensure data can be seamlessly passed through each pipeline module.

One such step is performed by eyeris::resample(), which safeguards the assumption of a fixed sampling rate on which the remainder of the pipeline relies. Rate-dependent stages such as eyeris::detransient(), eyeris::lpfilt(), and eyeris::downsample() assume that samples are evenly spaced in time. EyeLink trackers uphold this assumption by zero-filling missing pupil samples, but some hardware may instead drop samples entirely whenever pupil data is missing, leaving holes in an otherwise evenly-spaced time vector that silently distort any rate-dependent computation. eyeris::resample() repairs the time axis so that these potential differences in acquisition behavior do not propagate downstream.

For blocks that do require repair, eyeris::resample() proceeds in two stages. First, it constructs a uniform target grid anchored on the first reliable regular interval (i.e., the first observed interval matching the expected sampling period) rather than on the first timestamp; anchoring in this way prevents early sub-period jitter, for example, successive intervals of 3, 3, 4, and 4 ms under an expected 4 ms period, from offsetting the entire grid. This grid is then extended in both directions at the expected period so that it spans every observed timestamp, including any samples that precede the anchor. Second, the observed samples are placed onto the grid by linear interpolation: samples that fall on a grid point are retained verbatim, whereas short or jittered intervals are interpolated across so that their values contribute to the regular grid. Any observed interval longer than the expected period is considered a gap; the missing grid sample(s) within it are inserted as NA rather than interpolated across, and are flagged in a logical is_resampled column so that they can be tracked in subsequent processing stages.

#### 2.3.2 Blink artifact removal: eyeris::deblink()

First, preprocessing of pupil sizes needs to handle artifactual or missing data due to eye blinking, movement, or simply incorrect pupil detection and measurement by the eye tracker (see ***Figure 2a***). Most existing commercial eye trackers, including SR Research EyeLink^5^, have built-in blink detection methods that will annotate blink periods in exported data. One can easily recognize blink periods when the pupil size becomes 0/NaN for a brief period of time. However, it is important to recognize that there are often blink-related artifactual changes in pupil size surrounding an eye blink, which can be due to the eyelid rapidly covering the pupil area detected by the camera, resulting in rapidly diminishing pupil size before becoming zero. In contrast, as the eyelid opens, there is a brief period of rapidly increasing pupil size when the camera detects a partial pupil. Occasionally, eye trackers may incorrectly lock onto other structures in the camera’s field of view, such as eyebrows and glasses frames, as they may become more probable pupil targets compared to partially occluded pupils during blinking. This can also result in fluctuations in recorded pupil sizes.

**Figure 2.**
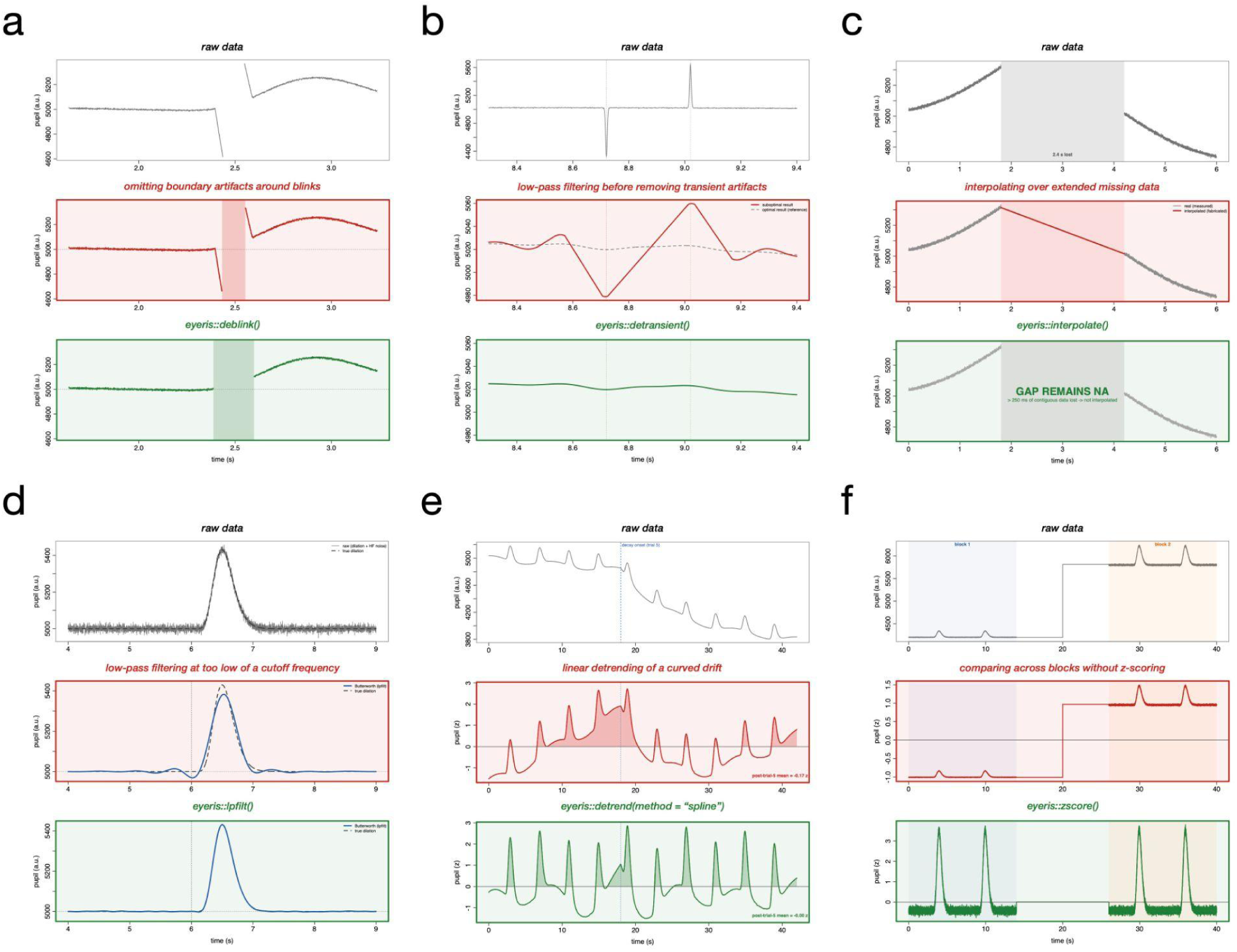
Six preprocessing pitfalls in pupillometry, each isolated on synthetic data with known ground truth signal. Each panel (a-f) contains 3 rows: the top row (grey) shows the input signal, the middle row (red border) shows the suboptimal processing choice, and the bottom row (green border) shows the recommended optimal processing choice using *eyeris* primitives. All traces are sampled at 1,000 Hz and were generated with the *eyeris::simulate_eyeris()* function under a fixed seed.

The deblink() function removes blink-related artifacts by NA-padding missing-data periods and their surrounding samples in the pupil time series. Each period of missing data is extended both backward and forward in time by a user-configurable duration (defaulting to 50 ms) to mask the partial-occlusion artifacts that typically flank eye blinks. The *extend* parameter accepts either a single value for symmetric padding or a two-element vector for asymmetric padding (e.g., *c(backward, forward)*). The appropriate *extend* duration may vary across experimental setups or participant populations and can be optimized upon inspection of the diagnostic reports generated by *eyeris*.

#### 2.3.3 Transient artifact removal: eyeris::detransient()

While commercial eye trackers can mark blink periods for easy exclusion, there are also artifactual pupil size measures that could occur in recordings due to unstable pupil tracking or eye/head movements. These non-blink disruptions of pupillometry measurement often result in transient and abrupt changes in recorded pupil sizes, commonly visible as spikes in the time series (see ***Figure 2b***). The detransient() function identifies and removes non-physiological pupil samples, such as transient spikes caused by unstable tracking or head movements, by applying a speed-based threshold. The function computes an approximate pupil change velocity using finite differences and flags samples whose speed exceeds a threshold derived from the Median Absolute Deviation (MAD; ***Arachchige and Prendergast, 2024; Hampel, 1968, 1974; Leys, 2013; Rousseeuw and Croux, 1993; Rousseeuw et al., 1986)*** of the velocity distribution. Specifically, the threshold is defined as the median speed plus *n* times the MAD value, where the constant *n* defaults to 16 ***(Tsukahara, 2020)***. For edge cases where the computed MAD or median speed is very small, users may supply a manual threshold via the *mad_thresh* parameter to adjust sensitivity. In our experience, most artifacts in pupillometry data can be captured using the simple detransient() method, thereby excluding non-physiological measures. However, noisier recordings with different eye tracker hardware may require more elaborate artifact detection algorithms, such as to remove sustained periods of outlying pupil sizes instead of velocity. Users are strongly encouraged to expand on the function using alternative detransient methods given their data and contribute to *eyeris* with additional methods to remove artifact periods (see ***Box 1*** and ***Appendix*** for more information about extensibility and developing custom extensions in *eyeris*).

#### 2.3.4 Linear interpolation: eyeris::interpolate()

Once data from blink and artifact periods are removed, the next step in pupillometry preprocessing is to interpolate over these periods to recover a contiguous time trace of pupil sizes ***(Mathôt et al., 2013)***. While not strictly required for certain types of pupillometry analysis, such as averaging across trial periods for a univariate measure, many recent analytic approaches for pupillometry data leverage each sampled time point to uncover temporal dynamics in changes of pupil sizes ***(Clewett et al., 2020; Johansson et al., 2018; Steinhauer et al., 2004; Urai et al., 2017; Van Slooten et al., 2018; Verney et al., 2004; Widmann et al., 2018)***. With these forms of analysis, avoiding missing data in certain periods provides more evenly distributed statistical power across time points (even with “guessing” of missing data through interpolation). The interpolate() function fills in missing pupil samples produced by the preceding blink-removal and artifact-detection steps, taking the simplest approach of linear interpolation given the relatively slow nature of pupil size fluctuations. It applies linear interpolation via the *na.approx()* function from the *zoo* package ***(Zeileis & Grothendieck, 2005)***, with edge NAs replaced by the nearest available values. Because linear interpolation becomes progressively less trustworthy as the span of missing data grows, *eyeris* does not interpolate across arbitrarily long gaps (see ***Figure 2c***). The *interpolate()* function exposes a *max_gap_ms* parameter that caps the maximum duration of a contiguous run of missing samples eligible for interpolation; runs exceeding this limit are left as missing (NA) rather than reconstructed. This threshold defaults to 250 ms, consistent with recommendations to restrict interpolation to short gaps ***(Kret & Sjak-Shie, 2019)***, and is converted internally to a sample count using each recording’s sampling rate so that the same temporal criterion applies regardless of tracker configuration. As with every preprocessing step, the value used is recorded in the JSON settings sidecar file (see ***Section 2.5)***, making the interpolation limit an explicit and auditable component of the analysis record.

In this respect, *eyeris* follows the guidance of ***Kret & Sjak-Shie (2019)*** to bound interpolation over long gaps, while deliberately departing from a one-size-fits-all exclusion rule for missing data. Rather than prescribing a fixed data-loss cutoff — a criterion whose appropriate value and level of application (block, epoch, or trial) depend on the experimental design — *eyeris* reports the proportion of missing samples (*prop_missing*) in its confound output at both the recording-block and trial/epoch levels (see ***Section 2.5)***. Researchers can therefore apply their own preregistered exclusion criteria at whichever granularity is appropriate for their analysis, and the corresponding percentage of data lost is surfaced in the *eyeris* HTML preprocessing report to keep this quality-control decision transparent.

#### 2.3.5 Low-pass filtering: eyeris::lpfilt()

It is evident that the interpolation steps should occur after missing-data detection, but it is less clear if, when, and how smoothing of typically noisy pupillometry data should take place. Smoothing of pupil sizes over time is commonly employed in the literature ***(Steinhauer et al., 2022)*** both to provide cleaner time trace visualizations and to reduce sporadic temporal dynamics that may be unrelated to underlying neurocognitive processes of interest. Smoothing is justified for pupil sizes with relatively slow dynamics but should always occur after blink and artifact detection to avoid contaminating neighboring samples during smoothing. In addition, smoothing can be achieved in multiple different ways, such as with moving averages or low-pass filtering. The lpfilt() function smooths the pupil time series by applying a low-pass Butterworth filter, with the default passband cutoff at 4 Hz and stopband at 8 Hz (see ***Figure 2d***). Note, the lpfilt() function here uses a minimum-order Butterworth filter, whose order is obtained with the buttord() function from the *gsignal* package ***(van Boxtel, 2021)*** to meet the defined passband and stopband cutoff frequencies (i.e., wp, ws) and ripple/attenuation requirements (i.e., rp, rs). The filter created by the *gsignal::butter()* function is returned in the second-order sections format. Users can easily modify the filter design parameters to achieve different smoothing results, but should always inspect the frequency response of the filter to ensure sufficient signal attenuation at higher frequencies and minimal ripples in the passband. The frequency response can be visualized by passing the plot_freqz = TRUE parameter to lpfilt().

#### 2.3.6 Downsampling: eyeris::downsample()

The downsample() function reduces the sampling frequency of the pupil time series to a user-specified target rate (*target_fs*) while preserving the original temporal dynamics of the signal. Unlike binning, downsampling retains individual data points after first applying an automatically designed anti-aliasing low-pass filter to prevent spectral aliasing. As a safeguard, the function raises an error if the resulting passband frequency would fall below 4 Hz, protecting frequency content relevant to pupillary responses. Users can optionally visualize the anti-aliasing filter’s frequency response by setting *plot_freqz = TRUE*.

#### 2.3.7 Binning: eyeris::bin()

The bin() function provides an alternative approach to reducing temporal resolution by dividing the time series into equal-duration intervals and computing a summary statistic — either the mean (default) or median — within each bin. The number of bins per second is controlled by the *bins_per_second* parameter, and the resulting timepoints are centered within each interval. Although binning is commonly used in pupillometry research, users should be aware that averaging within bins can attenuate rapid pupillary dynamics; the choice between downsampling and binning should therefore be guided by the specific research question.

#### 2.3.8 Detrending: eyeris::detrend()

Pupil sizes can fluctuate at a slow pace over the course of an entire recording; for example, the commonly reported decreasing trend during a cognitive task (a.k.a., the “time-on-task effect”; ***Beatty, 1982; Hopstaken et al., 2015; Martin et al., 2022; Massar et al., 2016; Steinhauer et al., 2022; Unsworth and Robison, 2016; Zhao et al., 2019)***. Depending on the scientific questions at hand, removal of this time-on-task effect may be desirable. *eyeris* provides the detrend() function to remove such slow trends over the entire recording (see ***Figure 2e***). The detrend() function fits a model of pupil size as a function of time and returns both the fitted values (the estimated trend) and the residuals (the detrended signal, i.e., pupil size minus the fitted trend). Two trend models are available via the *method* argument: the default, “linear”, removes a straight-line drift by fitting an ordinary linear model, whereas “spline” removes a smooth, potentially nonlinear drift by fitting a natural cubic spline basis of time (splines::ns(time, df = spline_df)). The spline option is well suited to cases in which the slow trend is poorly approximated by a single straight line, such as when the time-on-task effect accelerates or plateaus over the course of a recording. However, it should be noted that pupillometry analysis often employs baseline correction (see ***Mathôt et al., 2018b*** for a detailed discussion of the tradeoffs between subtractive and divisive baseline correction approaches) in a trial-by-trial design; as such, detrending might be unnecessary and could also increase the risk of introducing artifacts in the data if an extreme trend gets extracted (although unlikely as these artifacts were likely removed in the preceding steps). Moreover, high-pass filtering is discouraged as a way to remove slow trends in pupillometry data because designing a filter with a sufficiently sharp cutoff at the lower frequency end is difficult; failing to implement a proper filter design could drastically change the waveforms of pupil sizes that contain frequency content at approximately 1 Hz. Lastly, slow trends in pupil sizes might be physiologically meaningful, as tonic pupil sizes may reflect tonic arousal levels ***(Aston-Jones and Cohen, 2005; Groot et al., 2021; Joshi and Gold, 2020; Madore et al., 2020; Madore and Wagner, 2022; Murphy et al., 2011; Schwartz et al., 2025; Smallwood et al., 2012; Tsukahara et al., 2016; Unsworth and Robison, 2018)***. Depending on the target research question(s), such trends may need to be preserved for downstream analysis. Therefore, *eyeris* defaults to not performing detrending during preprocessing. Users may enable this step within *glassbox()* by setting *detrend = TRUE* when the removal of slow linear trends is appropriate for their research question.

#### 2.3.9 Z-scoring: eyeris::zscore()

There has been an ongoing debate regarding *z*-scoring normalization of pupillometry data. Some have argued against doing so outside of MRI contexts ***(Steinhauer et al., 2022)***. Normalizing pupil sizes by the mean and variance can introduce additional challenges in interpreting derived measures. Yet, it can also be argued that in the absence of normalization, pupil data across individuals cannot be easily pooled together for statistical inference, even if the non-normalized pupil sizes faithfully reflect individuals’ physiological responses. To illustrate, older adults and young adults may have differences in physical constraints that impact how much the pupil can dilate during cognitive performance. Not accounting for such differences may lead to the conclusion that older adults have weaker pupillary responses to certain stimuli (such as a loud tone). However, if different ranges of pupil fluctuations across individuals are accounted for by using *z*-scored pupil sizes, older adults may show comparable responses to such stimuli relative to their normal ranges of pupillary responses ***(He et al., 2020)***. Therefore, while older adults may show attenuated changes in pupil size due to physiological changes in muscles controlling pupil dilation, their cognitive arousal response to tone stimuli may not be preferentially reduced to suggest a selective deficit. Similar examples can be found even within a homogeneous group, since differences across individuals and across blocks/sessions are often present in each recording (e.g., ***Madore et al., 2020)***. Averaging non-normalized pupillary responses produces a metric biased by absolute pupil size, potentially confounding the cognitive (pupil) signal of interest with physiological variability.

Hence, *z*-scoring pupillometry data allows one to focus on the relative dynamics of fluctuations in pupil sizes rather than the first-level characteristics such as mean pupil sizes or absolute changes. It should be emphasized that *z*-scoring automatically makes the assumption that the variances accountable by changes in the absolute value or range of pupil sizes are irrelevant to target scientific questions. In many modern applications of pupillometry within psychological sciences, this is not an unreasonable assumption (i.e., ***Bradshaw, 1969; Gilzenrat et al., 2010; He et al., 2020; Peysakhovich et al., 2017; Van Gerven et al., 2004)***. Therefore, *eyeris* defaults to perform *z*-scoring normalization with the zscore() function. The zscore() function normalizes the pupil time series to have a mean of 0 and a standard deviation of 1 by mean-centering and scaling, analogous to *base::scale()*. This normalization facilitates trial-level and between-subjects analyses, as the arbitrary units of pupil size do not scale consistently across participants or recording sessions. Because *eyeris* is designed to preprocess one block of data per participant at a time, each block is z-scored independently (see ***Figure 2f***); when preprocessed data are later merged, all blocks will have been separately scaled. If a user supplies multi-block data that is not automatically separated by common start/stop recording indicators, we recommend the user to manually cut the data into multiple files (by block). *eyeris* supplies a bridge function that enables users to pass generic tabular eye-tracking data in case of situations like this and/or for trackers that are not natively supported by *eyeris* at the time of processing. Z-scoring is enabled by default and can be disabled by setting *zscore = FALSE* within *glassbox()*. One can easily turn off *z*-scoring during preprocessing, and we strongly encourage users to carefully consider how the decision to apply or forgo normalization of pupil sizes will impact downstream interpretations of their data, particularly when absolute pupil size or cross-session and cross-individual differences are relevant to the scientific question at hand.

### 2.4 Additional functions for epoching, exporting, and visualizing pupil data

In addition to the core preprocessing steps described above, *eyeris* provides built-in methods to conveniently extract tidy trial-based epochs with optional baseline correction, export BIDS-style derivatives and diagnostic HTML reports (see ***Figure 1***), and interactively browse diagnostic plots for tens of thousands of trials at each preprocessing step (see ***Figure 3***).^6^

**Figure 3.**
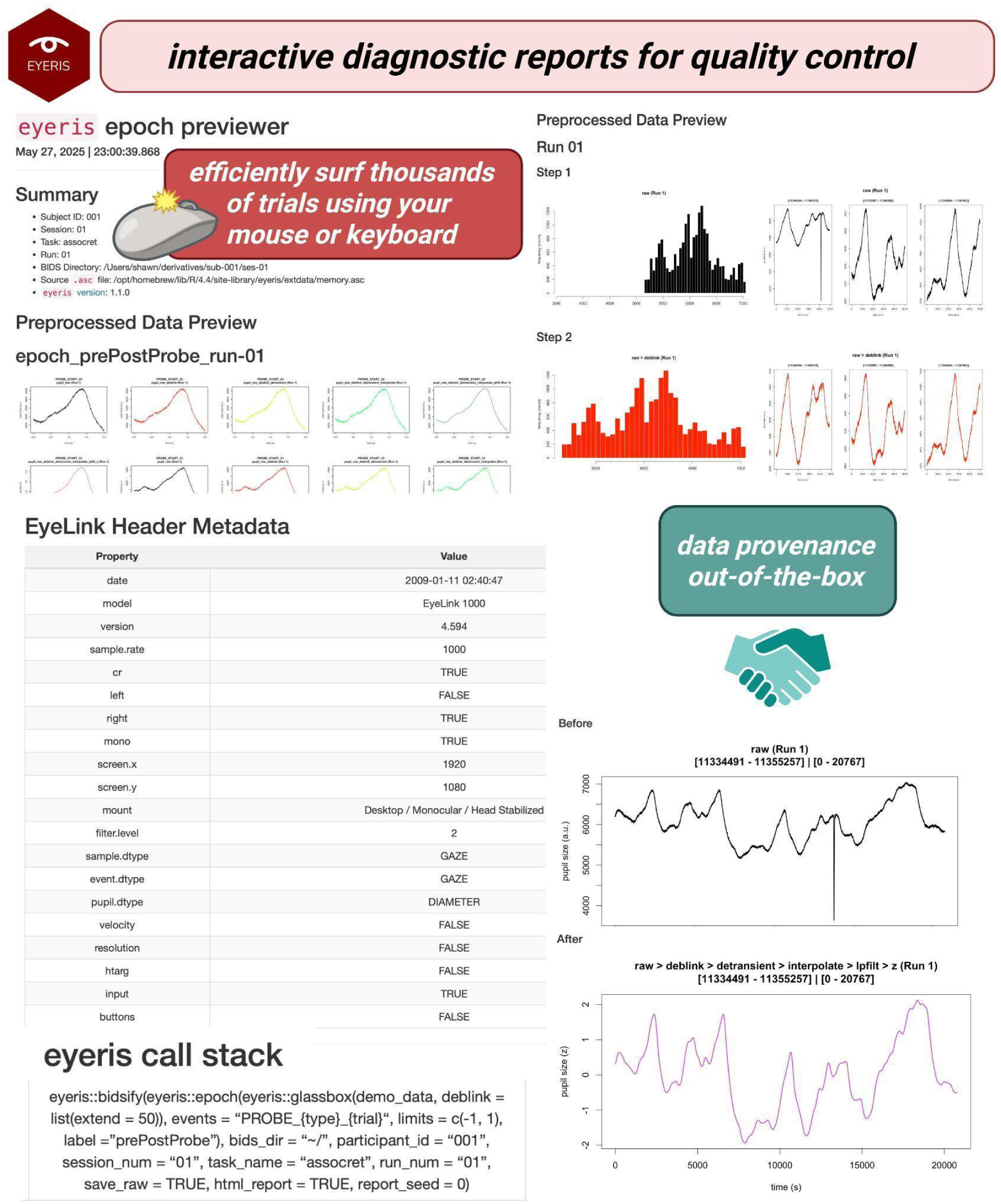
Overview of data and signal processing provenance provided out-of-the-box. *eyeris* provides automatically generated summary reports with diagnostic plots, recording metadata, and all preprocessing operations/parameters — and the resulting effects of each operation on the pupil time series — for every file/run for every participant. *eyeris* reports facilitate quality control by enabling surfing through thousands of trials worth of pupil data efficiently and interactively. Furthermore, these reports provide data provenance out-of-the-box and can be shared alongside any publication to support best practices in open science and reproducibility. Created in BioRender. *Schwartz (2025c)* https://BioRender.com/j2h2hzs.

#### 2.4.1 Epoching and baseline correction: eyeris::epoch()

The epoch() function is intended to be used as the final preprocessing step. It creates data epochs of either fixed or dynamic durations with respect to user-supplied event message strings and time limits, and includes an intuitive metadata parsing feature where additional trial data embedded within event messages can be automatically identified and joined into the resulting epoched data frames. The function supports three main event-modes: (*i*) a single string representing the event message to center epoch extraction around, using specified limits (or, if no limits are provided, extracting data between each subsequent occurrence of the same event); (*ii*) a vector containing both start and end event message strings, where limits are ignored and the duration of each trial epoch is determined by the number of samples between each matched start–end pair; and (*iii*) a list of two data frames that manually specify start/end event timestamp–message pairs for extraction from the raw time series, with explicit block number specification.

For event-modes (*i*) and (*ii*), the event message string must conform to a standardized protocol so that *eyeris* knows how to find events and optionally parse included metadata. Users have two primary choices: (a) specify a string followed by a *\** wildcard expression (e.g., *PROBE_START\**), which matches any messages beginning with that prefix; or (b) specify a string using the *eyeris* curly-brace syntax (e.g., *PROBE_{type}_{trial}*), which matches messages following that structure and automatically generates additional metadata columns (e.g., *type* and *trial*) in the tidy output. The function also supports optional baseline correction via the *calc_baseline* and *apply_baseline* parameters, with both subtractive and divisive correction types and flexible specification of the baseline period through either fixed time windows or event-delimited intervals (see ***Box 3*** for additional details).

Because all preceding preprocessing steps ***(Sections 2.3.1–2.3.9)*** operate on the continuous recording, epoching in *eyeris* is applied only after the pupil timeseries has been fully preprocessed. Interpolation, filtering, and detrending therefore act on the uninterrupted signal rather than on trial-level segments, avoiding edge artifacts at epoch boundaries and preserving information across trials.

###### Box 3. Tips & Tricks

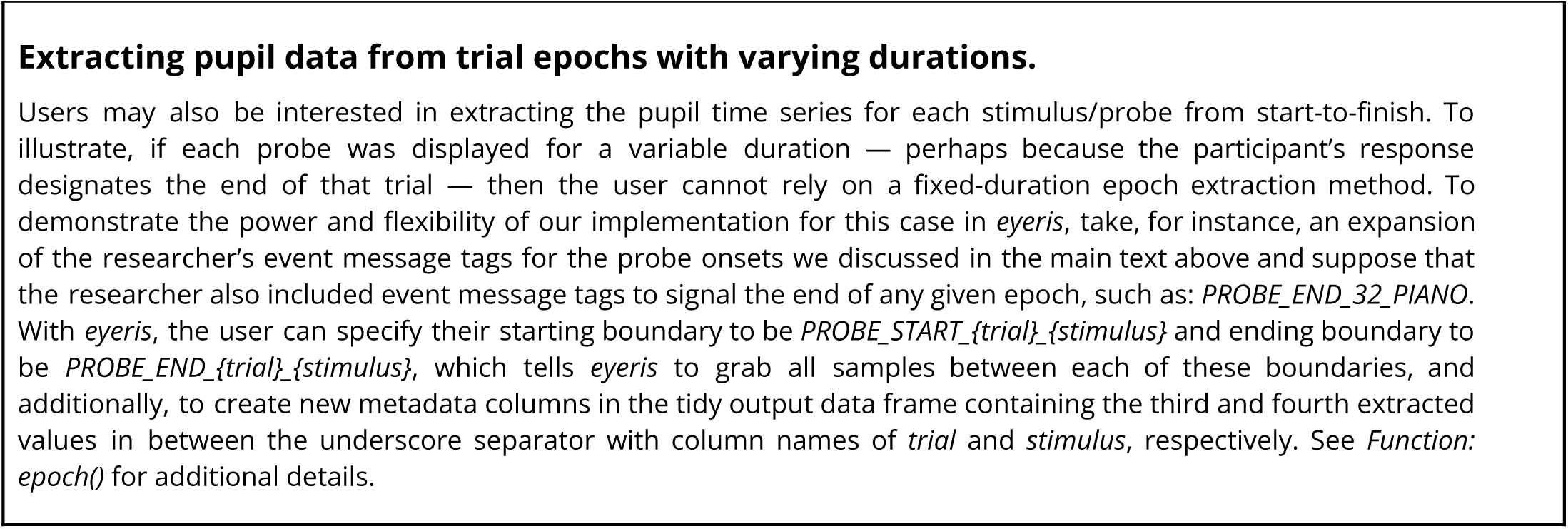

#### 2.4.2 Exporting BIDS-like derivatives: eyeris::bidsify()

The bidsify() function provides a structured way to save preprocessed pupil data in a BIDS-like directory structure. It exports epoched data as well as the raw pupil time series, and formats directory and file name structures based on user-supplied metadata (participant ID, session number, task name, and run number). For multi-block recordings contained within a single *.asc* file, blocks are automatically numbered as separate runs. The function also supports merging epochs into a single output file, merging runs across blocks, and optionally generating interactive HTML diagnostic reports. These reports include epoch-by-epoch diagnostic plots that can be grouped by a user-specified column variable (defaulting to the matched event label). In the future, we intend for this function to save data in an official BIDS format for eye tracking data (see the *<u>proposal currently under review</u>*); however, at this time, the function takes a BIDS-inspired approach to organizing output files (see ***Box 4)***.

#### 2.4.3 Diagnostic visualization: eyeris::plot()

The plot() function is an S3 plotting method for objects of class *eyeris*. It plots a single-panel time series for a subset of the pupil data at each preprocessing step, providing a simple method for qualitatively assessing the consequences of the preprocessing recipe and its parameters on the raw pupillary signal. Users can control which preprocessing steps to visualize (defaulting to all), the number and duration of random preview epochs, and whether to display accompanying histograms of pupil samples at each step via the *plot_distributions* parameter. A fixed random seed can be set to ensure the same preview epochs are displayed across repeated calls, which is especially useful when iteratively adjusting parameters within and across workflow steps. For multi-block recordings, the *block* parameter specifies which block to plot.

### 2.5 Documenting analytic decisions and data quality for reproducible sharing

As discussed above, *eyeris* retains configurability such that users may modify parameters, disable steps, and insert custom modules into the pipeline. Rather than eliminate these analytic degrees of freedom, *eyeris* is designed to structure and document them, so that the choices made during preprocessing are exposed and auditable rather than hidden. To this end, *eyeris* automatically emits two complementary machine-readable artifacts for every recording it processes: a per-run *confounds* table that quantifies data quality along many dimensions, and a per-run JSON metadata sidecar that records the exact preprocessing recipe that was applied. We recommend that researchers publish both of these files alongside their raw pupil data (see ***Section 2.5.3)***.

#### 2.5.1 Quantifying data quality: eyeris::summarize_confounds()

The *summarize_confounds()* function, invoked automatically within *bidsify()*, computes a battery of quality-control metrics *separately for each preprocessing step, each recording block, and each epoched time series* in the *eyeris* object. These metrics span five families: missing and invalid data (*n_missing, prop_missing, n_invalid, prop_invalid*); the temporal structure of data loss (*n_gaps* and the minimum, mean, and maximum gap length in both samples and milliseconds); blink statistics (*n_blinks, blink_rate_hz*, the minimum, mean, and maximum blink duration, *total_blink_time_ms,* and *prop_blink_time*); gaze position and dispersion (the horizontal and vertical gaze variance in pixels, and the mean gaze distance from screen center in both pixels and screen-normalized units); and pupil-size characteristics (the signal range, the proportion of samples pinned at the recording extremes, *prop_clipped*, and, at the trial level, the minimum and maximum z-scored pupil size and the pre-epoch and within-epoch pupil standard deviations). Missing data are reported as both a raw count (*n_missing*) and a proportion (*prop_missing*, ranging from 0 to 1; multiply by 100 for a percentage) of *NA* samples, and are computed at two levels of granularity so that users can choose the resolution appropriate to their design: at the block level, *prop_missing* is reported across an entire recording block; at the epoch/trial level, it is reported for each epoched event. *eyeris* additionally distinguishes *prop_missing* (samples that are *NA*, e.g., blinks and signal dropout prior to interpolation) from *prop_invalid*, which additionally folds in samples flagged as blinks or as off-screen gaze).

Critically, *eyeris* intentionally does not enforce a fixed missing-data exclusion cutoff. Instead, *prop_missing* is exposed as a column that users can filter on to define their own exclusion thresholds at whichever level — trial, epoch, or block — is appropriate for their study and hardware. This flexibility is deliberate given that recommendations for acceptable gap lengths and missing-data tolerances vary across experimental contexts and eye trackers ***(Kret and Sjak-Shie, 2019)***, and surfacing these quantities directly is arguably more transparent than silently discarding data behind a hard-coded rule. The confounds tables are written to the derivatives tree as *sub-XXX_task-XXX_run-XX_desc-confounds.csv* and, when database storage is enabled, are mirrored into the DuckDB backend (see ***Section 2.6)***.

#### 2.5.2 A machine-readable record of the preprocessing recipe: the JSON metadata sidecar

Every *eyeris* preprocessing step records its own call stack and the exact parameter values it was invoked with (stored internally in the object’s *params* field). As a consequence, the pipeline is able to self-document precisely what it did to a given dataset. When preprocessed data are exported, *eyeris* writes a per-run JSON metadata sidecar (*source/logs/run-XX_metadata.json*) that captures the run number, the source *.asc* file, and the ordered sequence of preprocessing operations together with the parameter values used at each step. Together with the *eyeris* version (which is also recorded at export) this sidecar constitutes a complete, machine-readable provenance record that is sufficient to reproduce the pipeline exactly. This mirrors the “glass box” provenance philosophy of reproducible neuroimaging tools such as *fMRIPrep* ***(Esteban et al., 2019)***.

From this same single source of truth, *eyeris* achieves workflow-level transparency by generating a copy-and-paste-ready, *fMRIPrep*-style methods boilerplate (*source/logs/methods_boilerplate.md*, also embedded in the diagnostic HTML report). This boilerplate is dynamically generated based on the workflow that was run and thus describes the executed workflow in plain prose with the actual parameter values substituted in, for example, the *deblink()* extension duration, the MAD multiplier used for transient rejection, or the low-pass filter passband and stopband cutoffs, so that a manuscript’s methods section can reproducibly reflect exactly what was run. Steps that were skipped are omitted and any user-supplied custom extensions are automatically described as well, so that non-default and multi-step pipelines are reported accurately. The generated boilerplate is released under a Creative Commons Attribution 4.0 (CC BY 4.0) license and is explicitly safe to paste directly into a manuscript, word-for-word, provided that *eyeris* is cited to satisfy attribution requirements.

#### 2.5.3 Sharing both the sidecar and the pupil data

We recommend that researchers using *eyeris* publish the JSON metadata sidecar and confounds tables *together with* their pupil data (raw and/or derivative). Because *eyeris* preserves configurability, distributing the sidecar exposes every preprocessing choice, including skipped steps and any non-default parameters supplied to the pipeline functions, rather than leaving those decisions implicit. This intentionally converts what would otherwise be a hidden set of researcher degrees of freedom into an auditable, reusable record, and pairs the executed code with human-readable reporting generated from the same provenance. In doing so, this practice directly advances the FAIR and open-science principles that motivate *eyeris*, and complements study preregistration by making the realized preprocessing pipeline transparent and independently reproducible.

### 2.6 Database storage and analysis

In addition to the BIDS-like file exports described further in ***Section 3.1.3***, *eyeris* provides an integrated database storage backend for centralized management of preprocessed pupillometry data. Database storage is activated by setting *db_enabled = TRUE* within the *bidsify()* function, which creates or connects to a DuckDB ***(Raasveldt & Mühleisen, 2019)*** database file (*.eyerisdb*) in the derivatives directory. All data types generated by the preprocessing pipeline, including time series, events, blinks, epochs, epoch summaries, and confounds, are automatically written to the database alongside any CSV exports. For large-scale deployments where storage efficiency and I/O performance are priorities, CSV output can be disabled entirely by setting *csv_enabled = FALSE*.

The *eyeris* database API provides a suite of functions for connecting to, querying, and extracting data from the database without requiring SQL expertise. The *eyeris_db_summary()* function provides a quick overview of the database contents, including available subjects, sessions, tasks, and row counts per data type. The *eyeris_db_collect()* function serves as the primary extraction interface, enabling researchers to aggregate data across multiple subjects in a single call with flexible filtering by subject, session, task, epoch label, and eye. For advanced use cases, direct SQL queries can be executed through the underlying DBI connection obtained via *eyeris_db_connect()*.

To support data sharing and distribution of large databases, *eyeris* includes chunked export functionality. The *eyeris_db_to_chunked_files()* function processes data incrementally (defaulting to 1 million rows per chunk) and writes the output to CSV or Parquet files, automatically splitting files that exceed a configurable size threshold (defaulting to 50 MB). For formal data distribution, *eyeris_db_split_for_sharing()* partitions the database into portable chunks that can be reconstructed by collaborators using *eyeris_db_reconstruct_from_chunks()*. The Parquet format is also supported, offering 50–80% file size reduction compared to CSV, faster read performance, and preserved numeric precision.

## 3 Results

### 3.1 Key Features & Innovations

Here, we introduce *eyeris* with the goal of making pupillometry preprocessing more efficient, reproducible, and conducive for reliable inferences when analyzing their pupillometry data.

#### 3.1.1 A modular signal processing workflow with carefully chosen parameters ensures efficient and robust pupillometry preprocessing

One key strength of the *eyeris* framework is its modularity. Each preprocessing function within *eyeris* is a wrapper around a core pipeline operation that automatically tracks, versions, and stores preprocessing states for any pupillometry dataset being processed in *eyeris*. This enables a “plug-and-play” experience when using *eyeris*: users have the option to use our recommended glassbox() method, as well as the ability to construct novel extensions and custom pipelines. Critically, this modular architecture promotes both adaptability and extensibility, facilitating the streamlined integration of custom preprocessing methods directly into the default workflow ***(Boxes 1***,***5***,***6***; ***Appendix)***.

###### Box 4. BIDS-like derivatives data structure

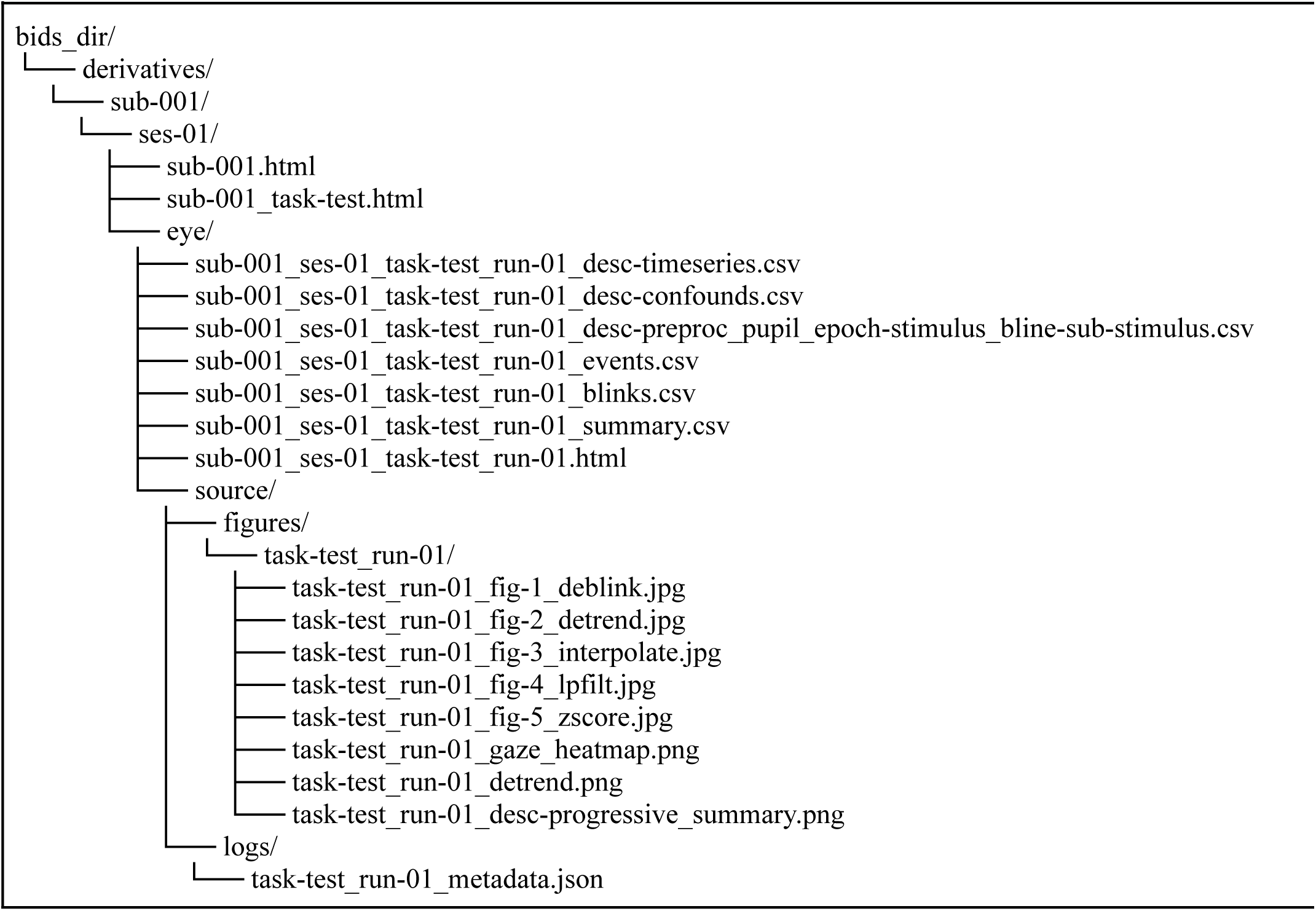

We recommend that users start with the glassbox() function to reduce the probability of inadvertently introducing sub-optimal computations or errors that could arise when reimplementing the individual steps underlying the preprocessing pipeline, both within and between projects. The glassbox() function allows users to make flexible adjustments, such as disabling a particular step and/or changing the default settings for a step, by passing in parameter arguments (see ***Box 6)***. A strength of the built-in glassbox() function compared to the many extant pupillometry packages is the framework it provides for rapidly preprocessing pupillometry data without requiring users to rely on copy-pasting complex step-by-step functions employed on a toy dataset from a vignette (which may not necessarily be optimized for the variety of forms that real-world pupillometry data come in). Furthermore, even when the instructions and functions supplied in a vignette of a package work “out-of-the-box” after pasting them into a custom script, it is still the user’s responsibility to ensure that every step was not only run, but done so in the correct order (with respect to the best practices of signal processing). Failing to do so could lead to unexpected and possibly detrimental results for the conclusions researchers will ultimately draw from the data. Therefore, we aim to facilitate this process to enable users to more easily engage with preprocessing steps by using a package, rather than moving away from them. With the *eyeris* glassbox() function, users can take a pupillometry recording file directly from a participant session, and in a short amount of time can go from raw to fully preprocessed data with detailed diagnostics for quality control in approximately three lines of code ***(Box 1***,***5***; ***Figure 3***).

###### Box 5. Tips & Tricks

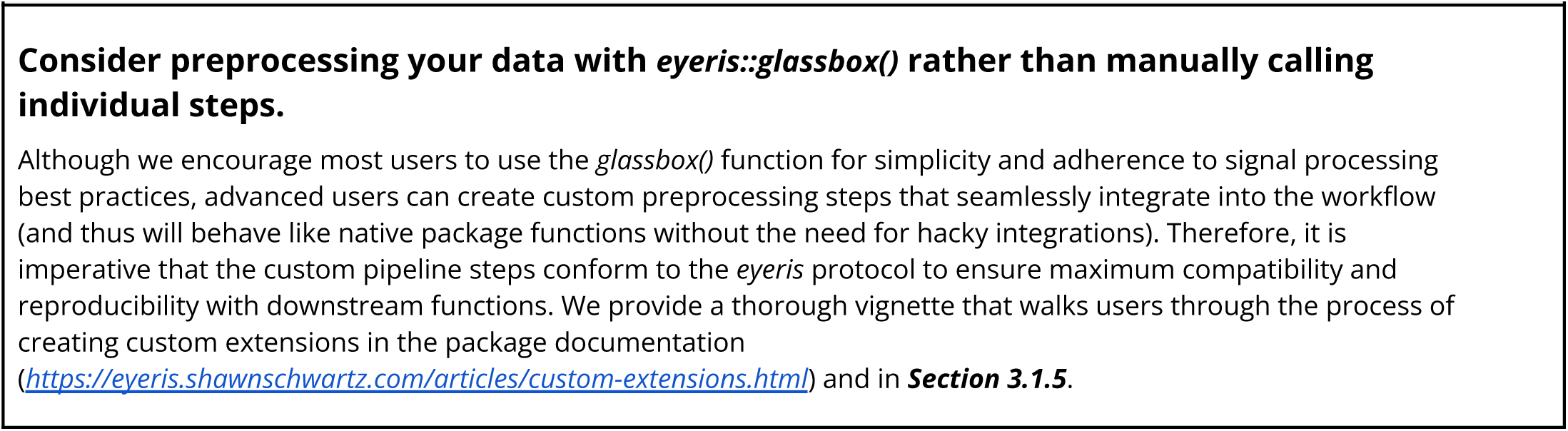

However, one critique of encapsulating a series of preprocessing steps into a single wrapper function, like glassbox(), is the regression towards a more black-box-like implementation. *eyeris* circumvents this pitfall in two key ways. First, the glassbox() function has two modes: interactive and headless. The interactive mode allows users to evaluate the consequences of the parameters used within each step on their data in real time, thereby facilitating an easy-to-use workflow for parameter optimization on any given dataset. In essence, this interactive process takes each recommended step and forces the user to evaluate both pre- and post-processing plots of time series for all steps in the default pipeline, with interjecting prompts to continue or modify parameters before moving forward. This stepwise approach to optimizing signal preprocessing parameters that yield satisfactorily de-noised pupil time series data reduces the opacity of the wrapper function to a transparent and engaging experience. The headless mode permits users to run the pipeline as-is on a dataset of any size — be it on their computer locally or on a cloud computing cluster — to preprocess large-scale pupillometry datasets using our recommended default parameters without manual intervention.

Second, *eyeris* stores the history of preprocessing steps and parameters for every action made on a raw dataset so that any derived data from the pipeline can be easily reproduced without additional steps needed from the researcher. Specifically, all of these data are saved to interactive HTML files (***Figure 3***), which contain numerous diagnostic plots for every step of the pipeline, automatically parsed tracker metadata, and a snapshot of the actions performed at each step of the workflow (see ***Section 3.1.3)***. Thus, using *eyeris* to deploy a large-scale preprocessing workflow on a headless computing architecture allows researchers to manually scrub their data sets for each run from every subject at every step of preprocessing without additional custom scripting. Critically, these interactive reports provided by *eyeris* go above and beyond within-study data quality control by automatically capturing and recording data and signal processing provenance in a standardized schema ***(MacKenzie-Graham et al., 2008)***, an essential feature for analytic reproducibility, lacking in many existing pupillometry preprocessing tools.

###### Box 6. Overriding default parameters in the *eyeris::glassbox()*

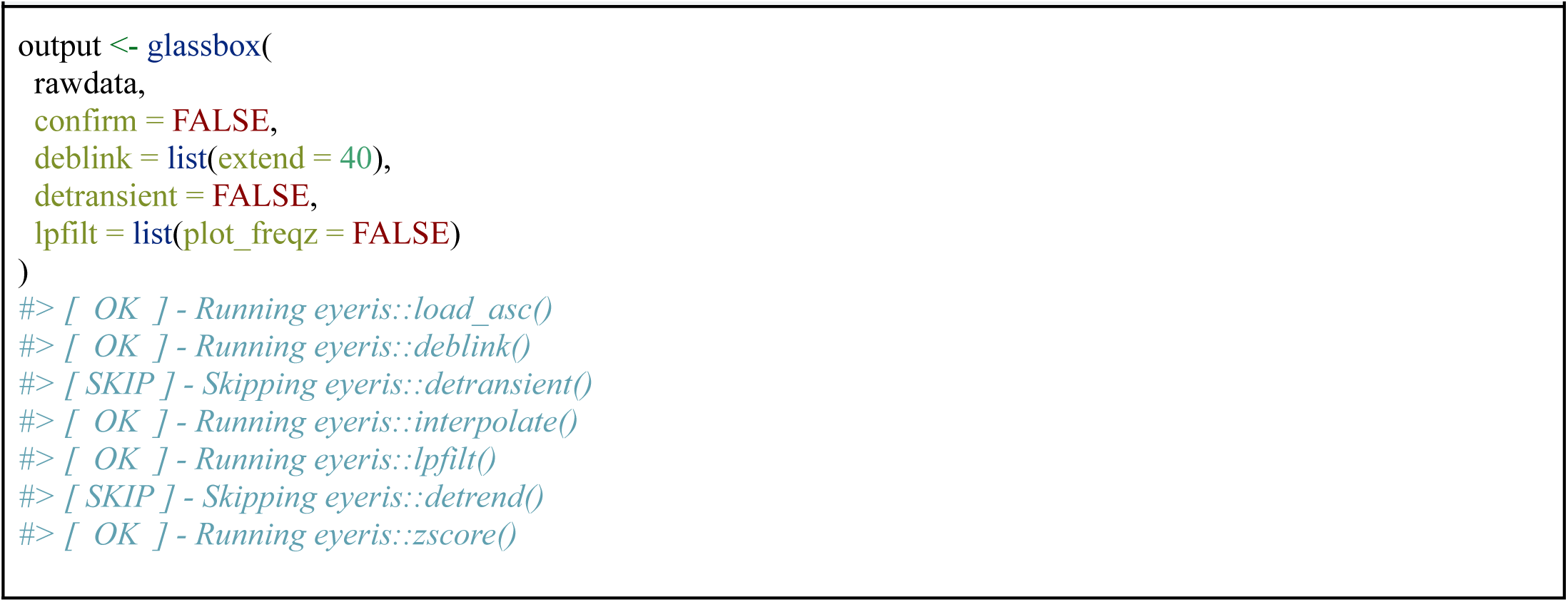

#### 3.1.2 User-friendly pattern matching and metadata parsing make extracting trial level epochs both accurate and effortless

Extant solutions for preprocessing pupil data lack intuitive workflows to extract trial-level epochs. For EEG researchers used to functions like pop_epoch in the MATLAB-based *EEGLAB* software (*Delorme and Makeig, 2004*), the lack of a comparable analogous function can lead to bottlenecks in a researcher’s pupillometry preprocessing workflow. Naturally, it made sense to take inspiration from *EEGLAB* when designing the epoch() function within *eyeris*. In particular, extracting trial-level data epochs with *eyeris* is flexible and offers advanced tooling for the extraction of metadata from event message strings. To illustrate, a researcher may have event message tags that look something like: PROBE_START_32_PIANO, indicating the timestamp for the stimulus probe onset on trial 32 where the stimulus was the piano. While not all event message strings contain as much information as this contrived example (e.g., standard EyeLink trial markers *TRIALID 1* often provide helpful trial-number metadata), the additional sources of information in the former example can be leveraged by the researcher as an additional sanity check to ensure accurate linking of pupil data with behavioral and/or neuroimaging data prior to analysis. Unsurprisingly, with more complex event message strings comes more burden on the researcher to devise an accurate and reliable solution to handle their specific event message configuration, likely leading to inefficient one-off solutions for particular datasets. *eyeris* meets this common demand by providing a carefully crafted regular expression implementation designed to handle a wide variety of generic cases ***(Box 3)***.

Conceptually, there are numerous ways that a researcher might want to extract their trial epochs for downstream analysis. In the simplest scenario, the user may be interested in testing a hypothesis about the tonic pupil size in the 1-second period just before the onset of the stimulus. In this case, the user would need to provide the string pattern and fixed start/stop duration of the epoch they would like to extract. For example, c(-1,0) for the 1-second just before the onset of the trial specifies a start/stop time array using the matched event message strings as the reference point (i.e., time=0).

#### 3.1.3 Interactive visual reports and a BIDS-like data scheme enable intuitive quality control and transparent data reporting

To date, most tools for preparing pupillometry data lack suitable features for data quality control up to the individual trial level. While some packages have integrated helpful plotting functions (e.g., ***Forbes, 2020; Hershman et al., 2019; Mittner, 2020; Tsukahara, 2020)***, these approaches only get the user part of the way there, as the onus falls on the user to aggregate such plots in a meaningful way for quality control. Moreover, this significant hurdle could be an arduous task even for researchers with programming experience. We strongly recommend that researchers carefully inspect the data at each stage of the preprocessing pipeline before aggregating trials and/or running inferential statistics, as is common in other human neuroscience data modalities such as EEG with the *EEGLAB* interactive viewer ***(Delorme and Makeig, 2004)***, M/EEG with *MNE-Python* ***(Gramfort et al., 2013)***, fMRI with *fMRIPrep* ***(Esteban et al., 2019)***, and structural MRI with *MRIQC* ***(Esteban et al., 2017; Provins et al., 2023)*** visual reports.

Given the lack of generic plug-and-play quality control solutions to date, in addition to the amount of effort required to implement such a solution, we suspect that many published pupillometry studies may not have raw data readily inspected end-to-end before and after preprocessing and running statistics. We therefore provide users with visualization tools of the pupil time series before and after each preprocessing operation for both an entire block/task and every trial-level epoch (i.e., as delineated by event messages/epochs). With this key feature, we hope to lower the barrier required for researchers to visually inspect entire recording sessions for anomalies in an efficient manner, as is necessary to ensure reproducible research results.

For the above reasons, we made visual reporting the default behavior in *eyeris* (***Figure 3***). Similar to the goals put forth by the team behind *fMRIPrep*, we anticipate that automatically generating diagnostic reports per participant will (*i*) maximize shareability and transparency between researchers, (*ii*) make data surfing less cumbersome and more efficient and hence increase the probability that thorough data surfing occurs before submitting data to statistical tests, (*iii*) increase statistical power of future pupillometry investigations by enabling researchers to purge poor data at the trial level without excluding entire runs or participants altogether, and (*iv*) contribute to best practices in open and reproducible science by recording all preprocessing decisions and settings for any given dataset that makes its way through *eyeris* by default ***(Esteban et al., 2020, 2019; Poldrack et al., 2019)***.

Importantly, all derived data and reports from the foregoing preprocessing workflows are automatically sorted into a BIDS-like format ***(Gorgolewski et al., 2016)***, facilitating separation of source (i.e., original or raw) data from derived (i.e., preprocessed, trimmed, epoched, etc.) data, in a directory structure that is intuitive to navigate, automatically organized, and therefore essentially ready to share on an appropriate data sharing outlet/repository like OSF, dryad, and/or GitHub (see ***Box 4)***.

#### 3.1.4 Integrated database storage enables scalable and efficient management of large-scale pupillometry datasets

As pupillometry research increasingly involves large sample sizes, multi-session designs, and high-throughput cloud computing deployments, traditional CSV-based data management workflows can become unwieldy. For studies with hundreds or thousands of participants — each generating multiple recording files, preprocessing derivatives, epoch extractions, and diagnostic metadata — the resulting proliferation of individual CSV files poses challenges for data organization, storage efficiency, and analytic tractability. Moreover, syncing and transferring thousands of small files across cloud storage systems incurs significant I/O overhead, and loading entire datasets into memory for analysis is often impractical.

To address these scalability challenges, *eyeris* provides an integrated database storage backend powered by DuckDB ***(Raasveldt & Mühleisen, 2019)***, a high-performance embedded analytical database engine. When database storage is enabled within the *bidsify()* function (by setting *db_enabled = TRUE*), all preprocessed data — including time series, events, blinks, epochs, epoch summaries, and confounds — are written to a single, self-contained database file (*.eyerisdb*). This centralized storage approach offers several key advantages over flat-file workflows. First, queries execute at the storage level, allowing researchers to filter, aggregate, and join data across subjects without loading entire datasets into memory. Second, a single database file dramatically reduces I/O costs on cloud platforms compared to syncing thousands of individual files. Third, built-in schema validation maintains data integrity and automatically tracks metadata such as creation timestamps, data types, and relationships between tables. Fourth, researchers can leverage SQL for complex cross-subject filtering and targeted extraction of data subsets, while the *eyeris_db_collect()* wrapper function provides a user-friendly interface that requires no SQL knowledge.

Critically, the database workflow is designed to complement, not replace, the existing BIDS-like file structure described above. Users may choose to generate both CSV files and a database simultaneously, maintaining human-readable outputs alongside the database for transparency, or opt for a database-only workflow to minimize storage and I/O costs in cloud computing environments (e.g., when deploying *eyeris* on high-performance computing clusters). This flexibility ensures that *eyeris* can scale from small desktop analyses involving a handful of participants to large-scale, multi-site pupillometry studies with hundreds or thousands of recordings, without requiring researchers to fundamentally change their preprocessing workflow.

For collaborative research and open-science workflows, *eyeris* also provides tools for distributing large databases in portable formats. The *eyeris_db_to_chunked_files()* function enables chunked export of large databases to CSV or Parquet files, processing data incrementally to avoid memory constraints and automatically splitting files that exceed a configurable size threshold. The Parquet format, in particular, offers substantial advantages for archival and sharing: 50–80% file size reduction compared to CSV, faster read performance, and preserved numeric precision. Together with *eyeris_db_split_for_sharing()* and *eyeris_db_reconstruct_from_chunks()*, these tools enable seamless data distribution and reconstruction, further supporting the FAIR principles that underpin *eyeris*.

#### 3.1.5 Advanced features for developers

We provide developers the flexibility to extend upon *eyeris*’ default offerings by writing their own custom pipeline operations that can integrate seamlessly with *eyeris* primitives (see ***Box 1)***.

Under the hood, each preprocessing function in *eyeris* is a wrapper around a core operation that gets tracked, versioned, and stored using the *pipeline_handler()* function. The *pipeline_handler()* enables flexible integration of custom data processing functions into the *eyeris* pipeline. It accepts an *eyeris* class object, the name of the operation function to apply, a character string suffix for the new column name, and any additional custom parameters. The function returns an updated *eyeris* object with the new column added to the *timeseries* data frame and the *latest* pointer updated to reflect the full history of preprocessing steps (e.g., *pupil_raw* → *pupil_raw_deblink* → *pupil_raw_deblink_detransient* → *pupil_…*). Following the *eyeris* protocol ensures that all operations follow a predictable structure and that new pupil data columns based on previous operations in the chain are dynamically constructed within the core time series data frame. Custom pipeline steps must conform to this protocol for maximum compatibility with the downstream functions we provide.

To demonstrate the flexibility of *eyeris* for developers, we walk through building a custom extension below. Please note that in the example that follows, we are *not* recommending that one performs this particular statistical transformation on their pupil data; rather, this is purely for demonstration purposes. Suppose you want to write a new *eyeris* extension function called *winsorize()* to apply winsorization^7^ to extreme pupil values. First, you write the core operation function, which should accept a data frame *x*, a string *prev_op* (the name of the previous pupil column), and any custom parameters. Second, you create a wrapper using the *pipeline_handler()* function, which enables your function to automatically: (a) track your function within the *eyeris* list object’s *params* field, (b) append a new column to each block within the *timeseries* list of data frames, and (c) update the object’s *latest* pointer. To ensure compatibility with *eyeris*, we impose the following requirements for any custom extension functions: (1) the first argument is always the *eyeris* class object; (2) the second is the name of the core operation function; (3) the third is the internal label you want *eyeris* to refer to (i.e., the one for the column name, plots, etc.); and (4) the fourth position onward are the custom parameters you defined in the core operation function. See ***Appendix*** for all source code for this demo. More information about customizing *eyeris* can be found in the online package documentation^8^.

## 4 Conclusions

The bells and whistles detailed throughout this manuscript and the documentation website^9^ for the *eyeris* R package aim to standardize the way pupillometry data are preprocessed and used in downstream analysis. To this end, we designed the architecture underlying *eyeris* to support the building of a robust suite of pupillometry tools analogous to the current offerings for fMRI analysts with *fMRIPrep* ***(Esteban et al., 2020, 2019)***. At its core, *eyeris* enables standardized and flexible preprocessing of the pupil assay.

Moreover, *eyeris* was designed with FAIR^10^ principles in mind. To fulfill this promise, *eyeris* uses a built-in protocol to carry out and manage the record keeping of the entire preprocessing pipeline on a raw dataset (see ***Figure 1***). This protocol is more or less a “grammar”, where each operation is a “verb” that can be chained one after the other (see ***Box 1***, which draws inspiration from familiar tidyverse “pipe” workflows; ***Wickham et al., 2019)***. This approach allows us to take advantage of syntactic sugar such that one could take a raw pupillometry dataset and within seconds obtain (*i*) a step-by-step record of all preprocessing steps (and parameters used), (*ii*) how each step sequentially impacted the raw signal, (*iii*) default diagnostic reports that organize relevant preprocessing metadata for any particular dataset passed through *eyeris*, and (*iv*) an intuitive gallery viewing functionality for efficiently scrolling through hundreds-to-thousands of raw data epochs for quality control prior to running inferential statistics (***Figure 3***).

Additionally, *eyeris* provides integrated database storage powered by DuckDB, enabling researchers to manage and query preprocessed pupillometry data from large-scale studies involving hundreds or thousands of participants within a single, self-contained database file. This database backend, combined with chunked export tools for portable data sharing in CSV and Parquet formats, ensures that *eyeris* can scale from small desktop analyses to cloud-deployed, multi-site studies without compromising on the transparency, reproducibility, or accessibility of the preprocessing workflow.

An important current limitation of *eyeris* is that native, fully validated support is provided only for SR Research EyeLink .asc files; other formats can be ingested through the generic loader described herein, but with default preprocessing parameters tuned to EyeLink-style data. While EyeLink systems remain widely used in cognitive pupillometry, this restriction excludes researchers using alternative eye-tracking hardware (e.g., Tobii, Pupil Labs, GazePoint). Extending *eyeris* to support additional systems involves more than implementing new file loaders: different eye trackers produce qualitatively different artifact profiles. For example, EyeLink systems tend to produce characteristic sharp downward spikes flanking blinks due to partial pupil occlusion by the eyelid, whereas video-based trackers may exhibit different artifact morphologies that could require adjusted detection parameters or different rejection strategies entirely. The modular architecture of *eyeris* — in particular, the *pipeline_handler()* system and the separation of file loading from downstream preprocessing — was designed to accommodate such extensions. The *load_asc()* function can serve as a template for future hardware-specific loaders (e.g., *load_tobii()*), and tracker-specific preprocessing steps can be integrated as custom extensions without modifying the core framework. To this end, *eyeris* provides a tracker-agnostic data ingestion function, *load_generic()*, which accepts standardized data frames; i.e., a required stream of raw samples (timestamps and pupil size) plus optional column-mapping specification for arbitrary tabular data, and assembles them into the same object structure returned by the default *load_asc()* method. Because the resulting object is a drop-in input to *glassbox()* and every downstream step, data from any tracker can enter the pipeline once exported to this common form, and format-specific readers can be layered on top incrementally. As several downstream steps assume a fixed sampling interval, *load_generic()* also warns when consecutive timestamps are not uniformly spaced, for instance, when a tracker drops samples rather than encoding missing data in place, so that such quirks are surfaced before preprocessing. We welcome community contributions of additional loaders via the project’s GitHub repository (see ***Code Availability)***. Despite this input variability, the signal processing principles underlying each preprocessing step (blink artifact removal, transient detection, interpolation, filtering, and normalization) are generally applicable across eye-tracking systems, even as specific default parameters may require re-tuning for different hardware.

We also note that *eyeris* is intentionally scoped as a preprocessing framework: it transforms raw tracker files into cleaned, epoched, and export-ready data, but does not itself implement statistical analysis or group-level visualization. However, the transition from preprocessing to analysis is straightforward. The epoched output from *eyeris* is a tidy data frame (one row per sample, with columns for participant, trial, condition metadata, time, and preprocessed pupil size) that is directly amenable to analysis with linear mixed-effects models (e.g., via *lme4* or *brms* in R), enabling researchers to analyze entire multi-participant datasets simultaneously without first computing participant-level averages. The DuckDB database backend further supports this workflow: *eyeris_db_collect()* can aggregate preprocessed data across all participants into a single table for group-level analysis and visualization with standard tidyverse tools. For detailed guidance on the statistical analysis stage of cognitive pupillometry, we refer users to ***Mathôt and Vilotijević (2023)***, whose analysis recommendations integrate naturally with the tidy output format produced by *eyeris*.

To this end, our hope is that enabling access to intuitive, interactive visual inspection tools to efficiently assess trial/epoch-level data quality, combined with scalable data management infrastructure, at scale, will support researchers in their efforts to perform reproducible and rigorous pupillometry research in a more accessible manner than is afforded by extant offerings.

## Acknowledgments

Development of this software library was supported by an agility grant from the Stanford University Wu Tsai Human Performance Alliance (to Prof. Anthony D Wagner & Shawn T Schwartz); the Stanford University Wu Tsai Neurosciences Institute Center for Mind, Brain, Computation and Technology (to Shawn T Schwartz & Alice M Xue); a Stanford University Ric Weiland Graduate Fellowship in the Humanities & Sciences (to Shawn T Schwartz); the McKnight Brain Research Foundation Clinical Translational Research Scholarship in Cognitive Aging and Age-Related Memory Loss; the American Brain Foundation; the American Academy of Neurology (to Haopei Yang); National Science Foundation Graduate Research Fellowship DGE-2146755 (to Alice M Xue); a Stanford Interdisciplinary Graduate Fellowship (to Alice M Xue); and National Institutes of Health Grants R01-AG065255 (to Prof. Anthony D Wagner) and R01-AG079345 (to Prof. Patrick L Purdon). The funders had no role in study design, data collection, and interpretation, or the decision to submit the work for publication. Furthermore, we are grateful to Ryan Yan for providing helpful comments on this manuscript. We are also grateful to Anthony Wagner for supporting the development of this software. We are thankful to Gustavo Santiago-Reyes for debugging and contributing to an earlier version of the source code, as well as to Natalie Wang for beta-testing the software during its early stages of development. We additionally thank Brian Knutson for helpful theoretical and practical discussions about pupillometry and arousal mechanisms. Lastly, we extend our gratitude to the past and present members of the Stanford Memory Lab^11^ who have influenced the work discussed herein.

## Author Contributions

Shawn T Schwartz, Conceptualization, Formal Analysis, Methodology, Project Administration, Resources, Software, Supervision, Validation, Visualization, Writing - Original Draft Preparation, Writing - Review & Editing; **Haopei Yang**, Data Curation, Resources, Validation, Writing - Review & Editing; **Alice M Xue**, Data Curation, Resources, Validation, Writing - Review & Editing; **Mingjian He**, Formal Analysis, Methodology, Resources, Supervision, Validation, Writing - Review & Editing.

## Code Availability

Please use the *issues* tab on GitHub to note bugs, comments, suggestions, and/or feedback.

- CRAN: https://cran.r-project.org/package=eyeris
- GitHub repository: https://github.com/shawntz/eyeris
- GitHub issues: https://github.com/shawntz/eyeris/issues
- Package documentation: https://eyeris.shawnschwartz.com

## Footnotes

1 See *Fink et al. (2024)* for a detailed table further outlining documented associations between cognitive processes and pupil size.

2 Throughout this manuscript, “preprocessing” means the specific steps necessary to derive transformed pupil data ready for visualization and analysis from the raw pupil data recorded from a participant using an eye tracking device.

3 See https://github.com/shawntz/clamp for a complete implementation of *eyeris* in headless mode for deployment on high-throughput cloud compute clusters (using the Slurm Workload Manager).

4 At this time, *eyeris* provides native, fully validated support for EyeLink-acquired pupillometry datasets, given its prevalence in this research space. Because accessibility to EyeLink hardware is limited by equipment cost, *eyeris* also provides *load_generic()*, a generic data ingestion mechanism that accepts standardized data frames (with an optional column-name mapping) so that recordings exported from any tracker can be brought into the pipeline, while noting that default preprocessing parameters remain tuned to EyeLink-style data. A more ambitious goal is native support for proprietary formats from other trackers; this step will require external contributions and requests from users with access to these data formats that we could use to implement the compatibility layer, which we will evaluate on a case-by-case basis via submissions to the issues tab on GitHub (see ***Code Availability*** for more information on how to access the source code and/or contribute directly to the development of *eyeris*).

5 https://www.sr-research.com

6 See https://eyeris.shawnschwartz.com/articles/ for example code and annotated walkthroughs.

7 https://en.wikipedia.org/wiki/Winsorizing

8 https://eyeris.shawnschwartz.com/articles/custom-extensions.html

9 https://eyeris.shawnschwartz.com

10 Findability, Accessibility, Interoperability, and Reusability (see *Hahn et al., 2024; Lazar, 2023; Wilkinson et al., 2016*).

11 https://memorylab.stanford.edu

